# OPUS-X: An Open-Source Toolkit for Protein Torsion Angles, Secondary Structure, Solvent Accessibility, Contact Map Predictions, and 3D Folding

**DOI:** 10.1101/2021.05.08.443219

**Authors:** Gang Xu, Qinghua Wang, Jianpeng Ma

## Abstract

In this paper, we report an open-source toolkit for protein 3D structure modeling, named OPUS-X. It contains three modules: OPUS-TASS2, which predicts protein torsion angles, secondary structure and solvent accessibility; OPUS-Contact, which measures the distance and orientations information between different residue pairs; and OPUS-Fold2, which uses the constraints derived from the first two modules to guide folding. OPUS-TASS2 is an upgraded version of our previous method OPUSS-TASS (*Bioinformatics* **2020,** *36* (20), 5021-5026). OPUS-TASS2 integrates protein global structure information and significantly outperforms OPUS-TASS. OPUS-Contact combines multiple raw co-evolutionary features with protein 1D features predicted by OPUS-TASS2, and delivers better results than the open-source state-of-the-art method trRosetta. OPUS-Fold2 is a complementary version of our previous method OPUS-Fold (*J. Chem. Theory Comput.* **2020,** *16* (6), 3970-3976). OPUS-Fold2 is a gradient-based protein folding framework based on the differentiable energy terms in opposed to OPUS-Fold that is a sampling-based method used to deal with the non-differentiable terms. OPUS-Fold2 exhibits comparable performance to the Rosetta folding protocol in trRosetta when using identical inputs. OPUS-Fold2 is written in Python and TensorFlow2.4, which is user-friendly to any source-code level modification. The code and pre-trained models of OPUS-X can be downloaded from https://github.com/OPUS-MaLab/opus_x.

## Introduction

Protein 3D structure prediction is crucial since the experimental approaches are usually time-consuming. In recent years, with the development of deep learning techniques, many methods have been proposed ^1–7^, improving the performance of protein structure prediction by a large margin. In the recent 14th Community-Wide Experiment on the Critical Assessment of Techniques for Protein Structure Prediction (CASP14), AlphaFold2 developed by DeepMind exhibits astonishingly performance ^7^, indicating that the computational methods have reached a practicable level.

Although protein 3D structure prediction is important, there are scenarios in which high-accuracy prediction of low-dimensional structural features, such as 1D features like torsion angles (*Φ* and *Ψ*), secondary structure (3-state and 8-state) and solvent accessibility, may be useful for successive modeling ^8^. Protein backbone torsion angles (*Φ, Ψ* and *Ω*) determine the entire protein conformation. Among them, *Ω* is around 180° in most case. Therefore, most researches only take *Φ* and *Ψ* into consideration ^9–12^. Protein secondary structure has been classified into either 3- or state ^13^, and it can be used to describe protein local conformation. Protein solvent accessibility measures the residue’s exposure to solvent at its folded state. Many successful methods have been proposed to predict protein 1D features ^9–12, 14, 15^, among which SPIDER3 ^11^ and NetSurfP-2.0 ^12^ adopted bidirectional recurrent neural networks to measure long-range interactions, SPOT-1D ^9^ integrated the predicted contact map ^16^ to capture protein global information. Our previous work OPUS-TASS^10^ introduced some new features derived from our potential functions ^17–19^ to improve the accuracy.

Protein contact map is critical to template-free modeling. At first, protein contact map is used to predict whether the Euclidean distance between two C_β_ atoms is less than 8.0 Å ^3, 16^. Then, some studies demonstrated the advantages of predicting real values of contact distance for the folding ^4, 20^. Recently, trRosetta ^5^ expanded the definition of contact information, including both distance and orientations information. In trRosetta, the distance information refers to the traditional C_β_- C_β_ distance, and the orientations information between residues 1 and 2 contains 3 dihedrals (ω, θ_12_, θ_21_) and 2 angles (φ_12_, φ_21_) ^5^. Here, ω denotes the dihedral of C_α1_- C_β1_- C_β2_- C_α2_, θ_12_ denotes the dihedral of N_1_-C_α1_- C_β1_- C_β2_, φ_12_ denotes the angle of C_α1_- C_β1_- C_β2_. Their results showed that orientations-guided folding performs better than distance-guided folding.

Protein 3D structure can be generated directly by optimization using energy-guided information. For instance, RaptorX-Contact ^3^ used Crystallography and NMR System (CNS) ^21^ to optimize its predicted distance constraints. trRosetta ^5^ developed a Rosetta protocol to optimize its distance and orientations constraints based on pyRosetta ^22, 23^. Currently, trRosetta-style’s folding is the most common one since it is fast and accurate.

In this research, we propose an open-source toolkit for protein 3D structure modeling, named OPUS-X. It consists of three modules: OPUS-TASS2, OPUS-Contact, and OPUS-Fold2. Comparing with its previous version OPUS-TASS ^10^, OPUS-TASS2 introduces the results from trRosetta ^5^ to measure its global information and adds protein solvent accessibility as its extra outputs. OPUS-Contact combines three raw co-evolutionary features similar to TripletRes ^24^ (including the covariance matrix (COV), the precision matrix (PRE) ^25^ and the coupling parameters of the Potts model by pseudo-likelihood maximization (PLM) ^26, 27^), the results from trRosetta ^5^, and the protein 1D features predicted by OPUS-TASS2 to deliver the final trRosetta-style’s outputs (ω, θ_12_, θ_21_, φ_12_, φ_21_). Different from our previous sampling-based protein folding framework OPUS-Fold ^8^, OPUS-Fold2 is a gradient-based method and can be used to perform the modeling guided by the trRosetta-style’s outputs from OPUS-Contact.

The contributions of this work can be summarized as follows:

1. The protein torsion angles, secondary structure, solvent accessibility predicted by OPUS-TASS2 are significantly more accurate than those predicted by the state-of-the-art methods in the literature.
2. The protein 3D folding performance of OPUS-Contact is better than that of trRosetta, which is an open-source state-of-the-art method.
3. We develop a flexible gradient-based protein folding method, OPUS-Fold2, which is written in Python and TensorFlow2.4, providing an alternative for the researchers who may need to modify the folding protocol or energy terms at source-code level. The accuracy of the results modeled by OPUS-Fold2 is comparable to that modeled by the Rosetta folding protocol in trRosetta.

## Methods

### Datasets

OPUS-TASS2 and OPUS-Contact use the same training and validation sets as OPUS-TASS ^10^, which were culled from the PISCES server ^28^ by SPOT-1D ^9^ on February 2017 with following constraints: resolution > 2.5 Å, R-free <1, and sequence identity < 25%. There are 10029 and 983 proteins in the training set and validation set, respectively.

In this research, we use 5 independent test sets to evaluate the performance of different approaches. CASP-FM (56), collected by SAINT ^29^, contains 10 template-free modeling (FM) targets from CASP13, 22 FM targets from CASP12, 16 FM targets from CASP11, and 8 FM targets from CASP10. CASP13 (26) contains 26 FM targets from CASP13. CASP14 (15) contains 15 FM targets from CASP14. The native structures of the targets in CASP13 (26) and CASP14 (15) are downloaded from the CASP website (http://predictioncenter.org). CAMEO-Hard61 (60), collected by OPUS-Rota3 ^30^, contains 60 proteins (one is discarded since it contains over 900 residues) released between January 2020 and July 2020, and labeled as hard targets by the CAMEO website ^31^. CAMEO (78), collected by trRosetta ^5^, contains 78 hard targets (we remove the targets that have missing residues for better evaluation) released between December 2018 and June 2019.

### Performance Metrics

MAE(*Φ*) and MAE(*Ψ*) are used to measure the mean absolute error (MAE) between the native protein backbone torsion angle and predicted one. SS3 and SS8 denote the percentage of correct prediction for 3- and 8-state protein secondary structure, respectively. ASA denotes the Pearson Correlation Coefficient of protein solvent accessibility.

To evaluate the performance of contact distance prediction, we use P_s ≥ 24_ and P_s ≥ 12_ to denote the precision of the top *L* predicted contacts with sequence separation of s, F/M and F/L to denote the F1-score of all possible contacts with sequence separation of 12 ≤s< 24 and 24 ≤s, respectively. TM-score ^32^ is used for protein 3D structure evaluation.

### Framework of OPUS-X

OPUS-X consists of three modules: OPUS-TASS2, OPUS-Contact and OPUS-Fold2. More details are shown in Figure 1.

**Figure 1.**
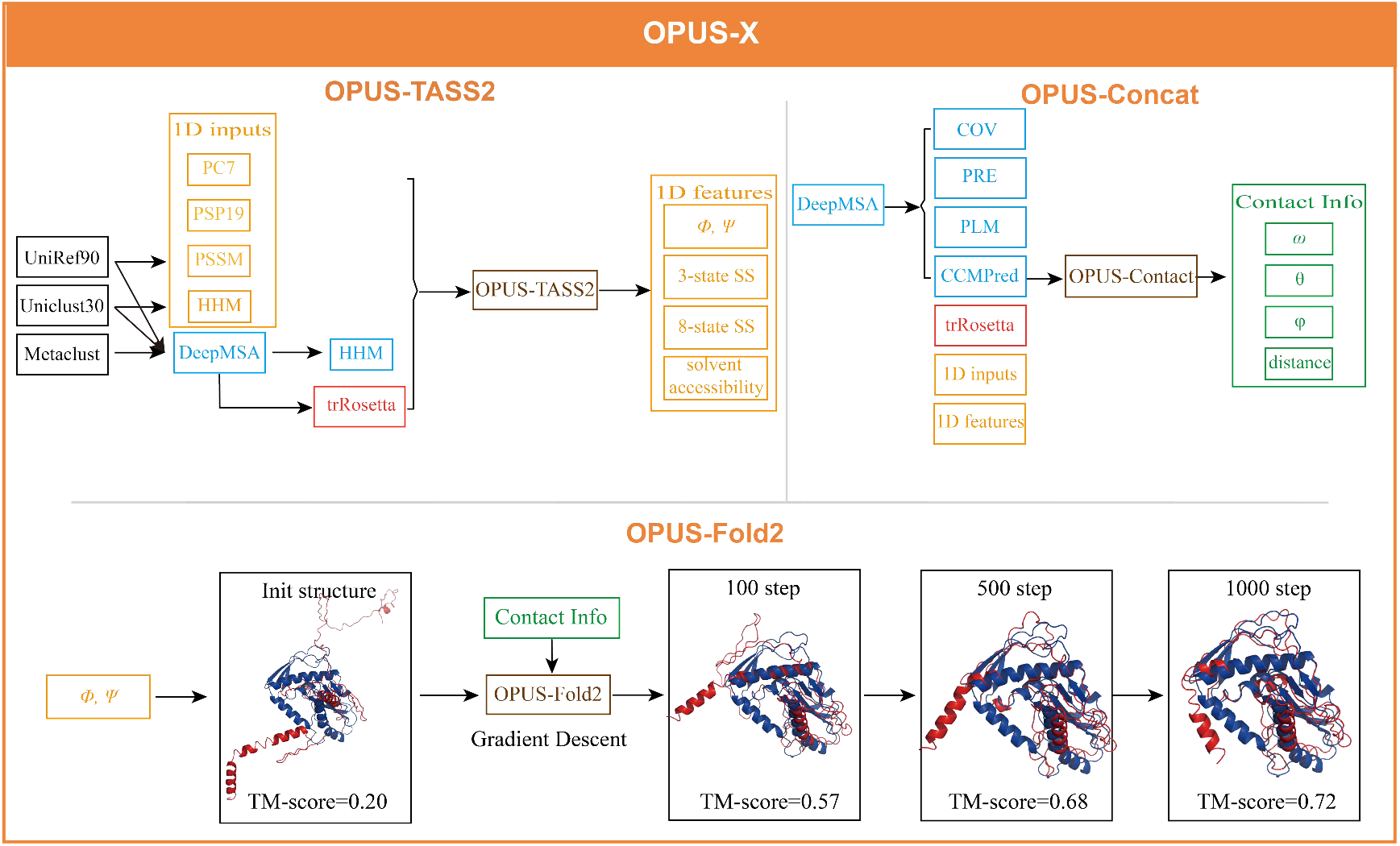
Three modules in OPUS-X. The red structures are the predicted structures during the folding, the blue structure is its native counterpart.

### OPUS-TASS2

The input features of OPUS-TASS2 can be categorized into three parts. The first part contains the same 76 features as OPUS-TASS ^10^, including 20 Position Specific Scoring Matrix (PSSM) profile features generated by three iterations of PSI-BLAST ^33^ v2.10.0+ with default parameters against UniRef90 database ^34^ updated in December 2019, 30 HHM profile features generated by HHBlits v3.1.0 ^35^ with default parameters against Uniclust30 database ^36^ updated in August 2018, 7 physicochemical properties and 19 PSP19 features ^17, 18^. The second part is 30 HHM profile features generated using *hhmake* from the multiple sequence alignment results obtained by DeepMSA ^37^. DeepMSA is a state-of-the-art multiple sequence alignment method which searches the alignment results in UniRef90, Uniclust30 and Metaclust ^38^. The third part is the output of trRosetta ^5^. Therefore, both distance and orientations global information can be captured in OPUS-TASS2.

The output features of OPUS-TASS2 contain one regression output node to predict solvent accessibility, 3 regression output nodes to predict CSF3 features ^10, 19^, 4 regression output nodes to predict sin(*Φ*), cos(*Φ*), sin(*Ψ*) and cos(*Ψ*), 11 classification output nodes to predict 3- and 8-state secondary structure. 8-state secondary structure is defined as follows: coil C, high-curvature S, *β*-turn T, *α*-helix H, 3_10_-helix G, *π*-helix I, *β*-strand E and *β*-bridge B ^9, 13^. They can be further classified into coil C (C, S and T), helix H (H, G and I) and strand E (E and B).

The neural network architecture of OPUS-TASS2 is shown in Figure 2. To introduce the results from trRosetta, we use a stack of dilated residual-convolutional blocks similar to trRosetta to perform the feature extraction. Sequentially, we perform feature selection, using the 64 filter-dimension features at (*n*, *n*) to represent the features of residue *n*. Then, we concatenate the 2D inputs with the 1D inputs, which contain the first part and second part input features of OPUS-TASS2 defined above, and feed them into the following modules which are basically identical to that in OPUS-TASS ^10^.

**Figure 2.**
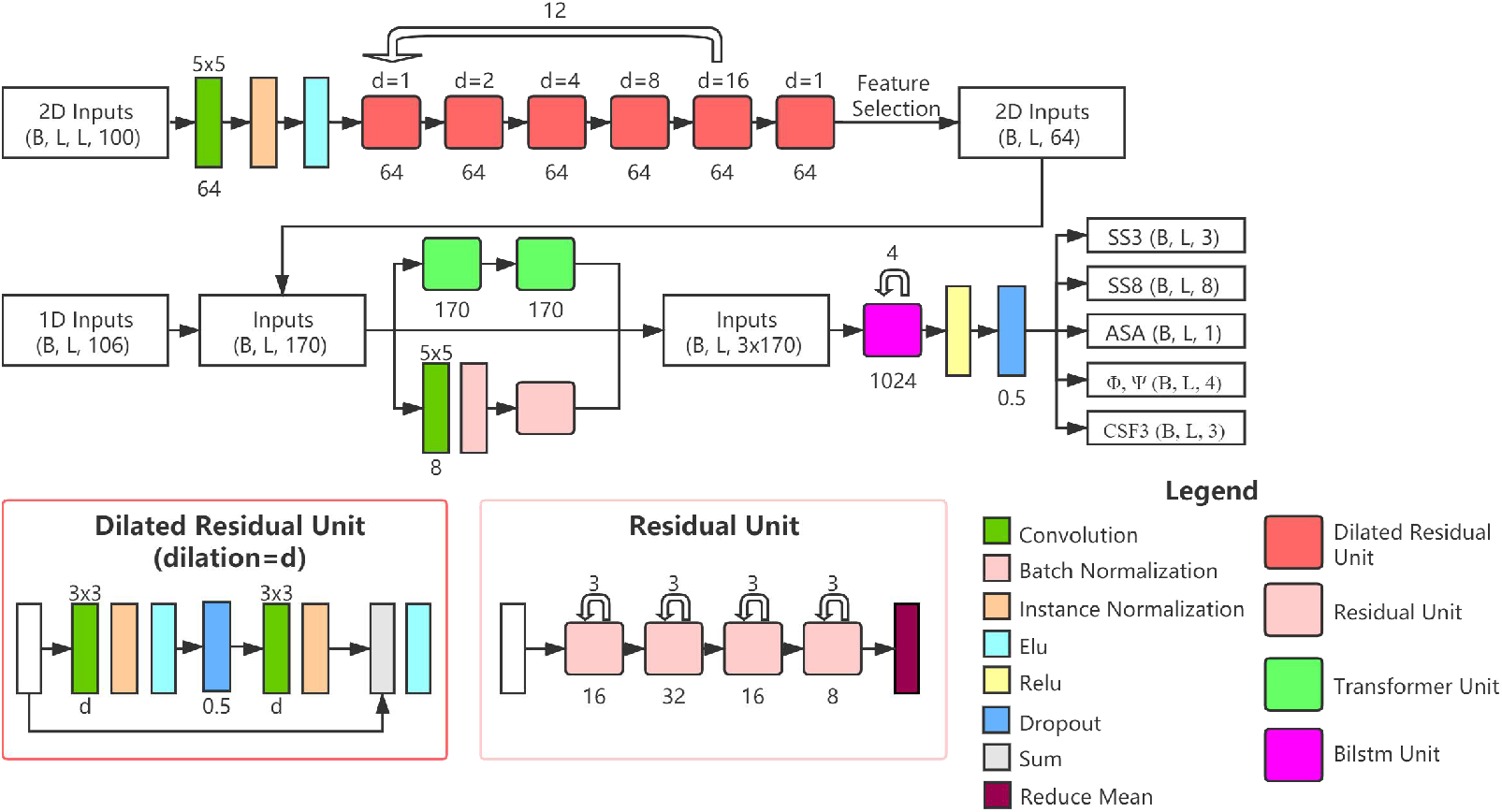
Framework of OPUS-TASS2. The outputs of trRosetta go through 61 dilated residual-convolutional blocks and output 64 features for each residue in which distance and orientations global information are included. Then, these 64 features and 106 1D features are concatenated to go through three modules: Resnet module ^39^, modified Transformer module ^10, 40^ and bidirectional Long-Short-Term-Memory module ^41^. The residual unit used here is similar to the conventional residual block except the strides of all units are set to be one. *B* denotes batch size, which is set to be one in OPUS-TASS2. *L* denotes sequence length.

OPUS-TASS2 adopts ensemble strategy as OPUSS-TASS ^10^ and SPOT-1D ^9^, and it consists of 9 models. The average is used for 3- and 8-state secondary structure classification prediction, and the median is used for backbone torsion angles and solvent accessibility regression prediction.

### OPUS-Contact

The inputs of OPUS-Contact contain 4 parts. Following TripletRes ^24^, the first three parts are the three raw co-evolutionary features: the covariance matrix (COV), the precision matrix (PRE) ^25^ and the coupling parameters of the Potts model by pseudolikelihood maximization (PLM) ^26, 27^. The fourth part contains 92 1D features: including 76 features from the first part of input features of OPUS-TASS2, and 1 solvent accessibility, 4 torsion angles (sin(*Φ*), cos(*Φ*), sin(*Ψ*) and cos(*Ψ*)) and 11 secondary structure (3- and 8-state) predicted by OPUS-TASS2. We use outer concatenation function as SPOT-1D ^9^ to convert 1D features (*L*, 92) into 2D features (*L*, *L*, 184). Together with the results from trRosetta ^5^ (*L*, *L*, 100) and CCMpred ^42^ (*L*, *L*, 1), the final fourth part features have 285 features in total. Here, COV, PRE, PLM, CCMpred and the results from trRosetta are generated from the multiple sequence alignment results obtained by DeepMSA ^37^.

The outputs of OPUS-Contact are identical to that of trRosetta ^5^, which include the predicted C_β_-C_β_ distance, 3 dihedrals (ω, θ_12_, θ_21_) and 2 angles (φ_12_, φ_21_) between residues 1 and 2. The distance ranges between 2 and 20 Å, and it is segmented into 36 bins with 0.5 Å interval, plus one bin represents the >20 Å case. φ ranges between 0 and 180°, and it is segmented into 12 bins with 15° interval, plus one bin represents the non-contact case. ω, θ range between −180 and 180°, and they are segmented into 24 bins with 15° interval, plus one bin represents the non-contact case.

The neural network architecture of OPUS-Contact is shown in Figure 3. We use a stack of dilated residual-convolutional blocks similar to the 2D feature extraction step in OPUS-TASS2. The 4 inputs parts (COV, PRE, PLM and Others) go through 41 blocks separately at first, and then concatenate to go through the following 21 blocks.

**Figure 3.**
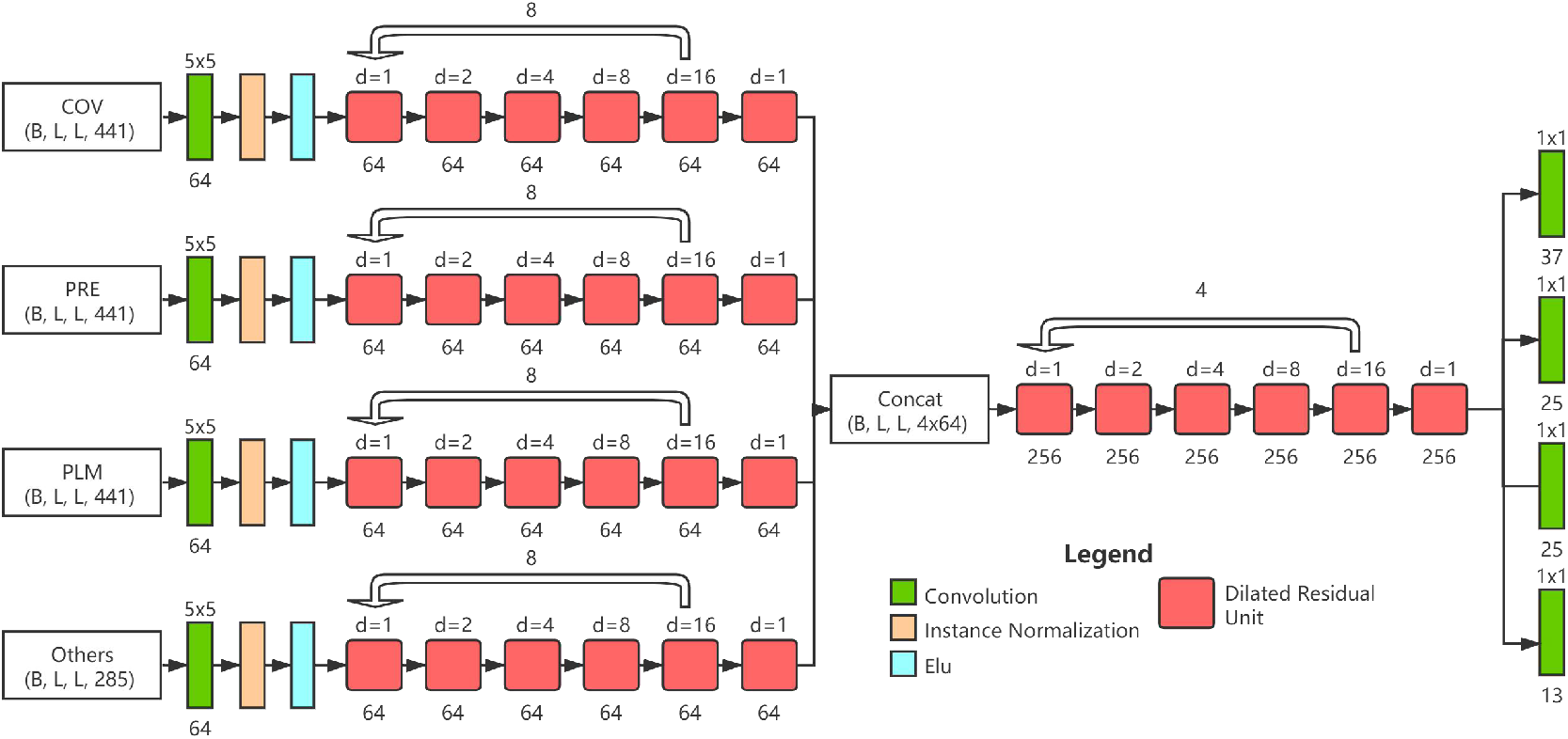
Framework of OPUS-Contact. The dilated residual unit is identical to that in OPUS-TASS2. *B* is also set to be one in OPUS-Contact.

OPUS-Contact also adopts ensemble strategy as trRosetta ^5^ and it consists of 7 models. The average is used for the final prediction.

### OPUS-Fold2

OPUS-Fold2 is a gradient-based protein folding framework. The variables of OPUS-Fold2 are the backbone torsion angles (*Φ, Ψ* and *Ω*) of all residues. OPUS-Fold2 minimizes the loss function derived from the outputs of OPUS-Contact by adjusting its variables.

The initial *Φ, Ψ* are predicted by OPUS-TASS2, and *Ω* is set to 180°. The loss function of OPUS-Fold2 in this research is defined as follows:

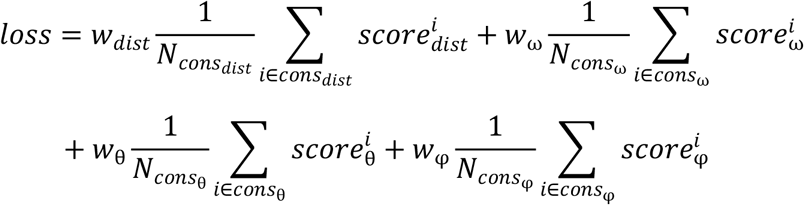

*cons_dist_* is the collection of distance constraints, in which *P_4≤dist<20_ ≥ 0.05*. *cons_ω_* and *cons_θ_* are the collections of ω and θ constraints, respectively, in which *P_contact_ ≥ 0.55*. *cons_φ_* is the collections of φ constraints, in which *P_contact_ ≥ 0.65*. *w_dist_*, *w_ω_*, *w_θ_* and *w_φ_* are the weights of each term, which are set to be 10, 8, 8 and 8, respectively. Similar to the folding protocol in trRosetta ^5^, we convert the distance and orientations distributions to the energy terms by the following equations:

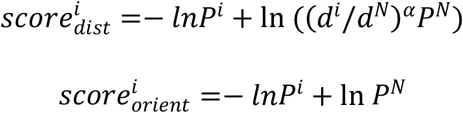

Following Dfire ^43^, the α is set to be 1.57. The reference state for the distance distribution is the probability of the *N*th bin [19.5, 20], and for the orientation distribution is the probability of the last bin [165°, 180°]. *d^i^* is the distance for the *i*th distance bin. *P^i^* is the probability for the *i*th bin. Cubic spline curves are generated to make the energy terms differentiable.

The optimization process of OPUS-Fold2 is based on TensorFlow2.4 ^44^, which is a flexible commonly-used tool to deal with the gradient descent tasks. We use Adam ^45^ optimizer to optimize our loss function with an initial learning rate of 0.5, 1000 steps are performed.

## Results

### Performance of OPUS-TASS2

We compare the performance of OPUS-TASS2 with that of NetSurfP-2.0 ^12^, SPOT-1D ^9^ and our previous work OPUS-TASS ^10^ on CAMEO-Hard61 (60), CASP-FM (56) and CASP14 (15). The results of NetSurfP-2.0 and SPOT-1D are obtained from their official websites. As shown in Table 1, OPUS-TASS2 achieves the highest accuracies for 3- and 8-state secondary structure prediction, the lowest mean absolute errors for torsion angles (*Φ* and *Ψ*) prediction and the highest Pearson Correlation Coefficient for solvent accessibility prediction on all three datasets.

**Table 1.** Performance of different predictors on CAMEO-Hard61 (60), CASP-FM (56), and CASP14 (15). The best result for each test is shown in boldface.

| | SS3 | SS8 | MAE($\Phi$) | MAE($\Psi$) | ASA |
| --- | --- | --- | --- | --- | --- |
| CAMEO-Hard61 (60) |  |  |  |  |  |
| NetSurfP-2.0 | 83.78 | 70.38 | 20.1 | 29.99 | 0.779 |
| SPOT-1D | 83.69 | 70.72 | 19.55 | 29.97 | 0.775 |
| OPUS-TASS | 84.15 | 72.12 | 19.26 | 29.47 | - |
| OPUS-TASS2 | **84.55** | **72.5** | **19.07** | **28.79** | **0.797** |
| CASP-FM (56) |  |  |  |  |  |
| NetSurfP-2.0 | 80.68 | 69.14 | 19.94 | 31.43 | 0.749 |
| SPOT-1D | 82.37 | 71.11 | 19.39 | 30.1 | 0.744 |
| OPUS-TASS | 83.4 | 73.27 | 18.85 | 28 | - |
| OPUS-TASS2 | **85.96** | **76.28** | **17.94** | **25.17** | **0.804** |
| CASP14 (15) |  |  |  |  |  |
| NetSurfP-2.0 | 75.39 | 61.87 | 22.62 | 40.54 | 0.68 |
| SPOT-1D | 75.19 | 61.41 | 23.19 | 43.98 | 0.663 |
| OPUS-TASS | 77.3 | 63.53 | 21.91 | 38.93 | - |
| OPUS-TASS2 | **80.87** | **68.26** | **20.53** | **33.48** | **0.735** |

The major differences between OPUS-TASS2 and OPUS-TASS ^10^ are the two extra input features, which are the second part 30 HHM profile features and the third part 64 global information features. To verify the importance of these two extra input features, we add them to the OPUS-TASS original 76 input features one by one. The results are shown in Table 2, it suggests that both of them are beneficial to the final prediction accuracy, especially the third part which contains the distance and orientations global information predicted by trRosetta ^5^.

**Table 2.** Importance of different parts in OPUS-TASS2 input features.

| 1<sup>st</sup> 76-d | 2<sup>nd</sup> 30-d | 3<sup>rd</sup> 64-d | SS3 | SS8 | MAE($\phi$) | MAE($\psi$) | ASA |
| --- | --- | --- | --- | --- | --- | --- | --- |
| CAMEO61 (60) |  |  |  |  |  |  |  |
| ✓ |  |  | 83.55 | 71.06 | 19.59 | 29.52 | 0.786 |
| ✓ | ✓ |  | 83.45 | 71.39 | 19.6 | 29.51 | 0.782 |
| ✓ | ✓ | ✓ | **83.68** | **71.58** | **19.17** | **29.24** | **0.788** |
| CASP-FM (56) |  |  |  |  |  |  |  |
| ✓ |  |  | 84.02 | 74.22 | 18.58 | 26.91 | 0.77 |
| ✓ | ✓ |  | 84.14 | 74.27 | 18.72 | 26.65 | 0.767 |
| ✓ | ✓ | ✓ | **85.85** | **75.63** | **18.35** | **25.47** | **0.795** |
| CASP14-FM (15) |  |  |  |  |  |  |  |
| ✓ |  |  | 76.02 | 62.41 | 21.79 | 38.65 | 0.68 |
| ✓ | ✓ |  | 77.3 | 64.15 | 21.92 | 36.91 | 0.705 |
| ✓ | ✓ | ✓ | **80.37** | **67.1** | **20.73** | **33.59** | **0.733** |

Since global information is crucial for protein 1D features prediction, we would like to find out the best performance OPUS-TASS2 can achieve if the input features for global information are all from the native structures. In Table 3, we list the performance of OPUS-TASS2 using the real orientation information (ω, θ and φ), real distance information, and both of them, respectively. The results show that, after introducing the real values, the performance of OPUS-TASS2 is significantly improved, which means the accuracy of OPUS-TASS2 can be increased by the improvement of trRosetta-style’s outputs.

**Table 3.** Performance of OPUSS-TASS2 based on the real values of different global information.

| | SS3 | SS8 | MAE($\phi$) | MAE($\psi$) | ASA |
| --- | --- | --- | --- | --- | --- |
| CAMEO61 (60) |  |  |  |  |  |
| OPUS-TASS2 | 84.55 | 72.5 | 19.07 | 28.79 | 0.797 |
| w/real orient | 89.29 | 81.37 | 15.9 | 17.39 | 0.877 |
| w/real dist | 87.08 | 76.42 | 17.91 | 24.14 | 0.835 |
| w/real all | **90.85** | **83.35** | **15.32** | **15.97** | **0.888** |

| CASP-FM (56) |  |  |  |  |  |
| --- | --- | --- | --- | --- | --- |
| OPUS-TASS2 | 85.96 | 76.28 | 17.94 | 25.17 | 0.804 |
| w/real orient | 89.46 | 82.81 | 15.2 | 16.25 | 0.885 |
| w/real dist | 88.04 | 79.33 | 16.87 | 21.47 | 0.85 |
| w/real all | **90.85** | **84.42** | **14.68** | **15.35** | **0.9** |
| CASP14-FM (15) |  |  |  |  |  |
| OPUS-TASS2 | 80.87 | 68.26 | 20.53 | 33.48 | 0.735 |
| w/real orient | 87.51 | 78.01 | 16.5 | 17.83 | 0.874 |
| w/real dist | 84.9 | 72.45 | 18.92 | 26.82 | 0.805 |
| w/real all | **90.04** | **81** | **15.41** | **16.45** | **0.889** |

### Performance of OPUS-Contact

To evaluate the performance of contact distance information, current studies ^3, 5, 16^ usually used precision of the top *L* predicted contacts or F1-score as the metric. However, as shown in Figure 4, the correlation between theses distance-based metrics and the TM-score of their corresponding 3D structures modeled by the folding protocol in trRosetta ^5^ is not significant.

**Figure 4.**
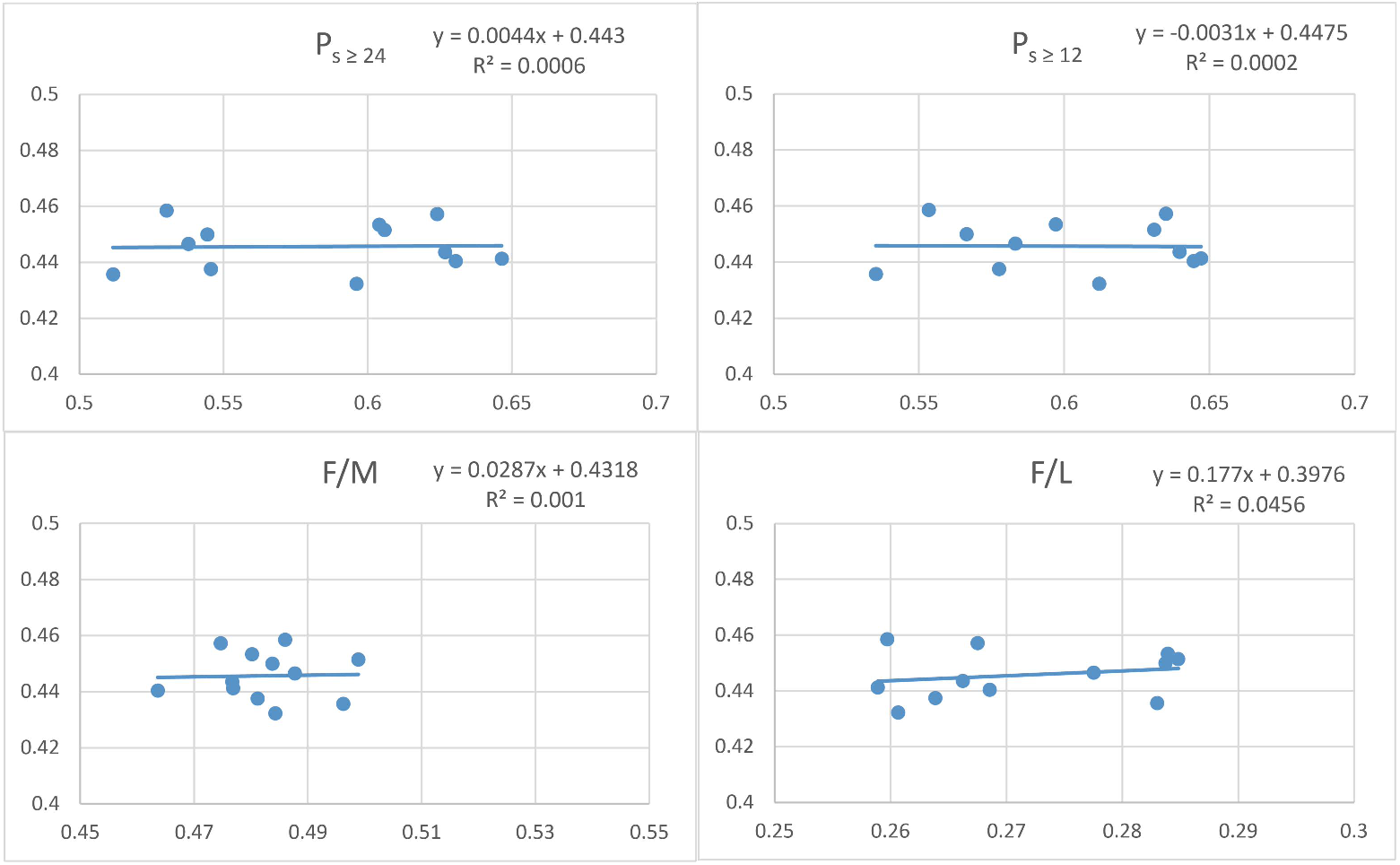
Correlation between distance-based metrics (P_s ≥ 24_, P_s ≥ 12_, F/M and F/L) and the TM-score of their corresponding 3D structures based on different OPUS-Contact models on CASP14 (15). Each dot denotes an OPUS-Contact model. The x-axis represents the result of each metric and the y-axis represents the TM-score.

The outputs of OPUS-Contact contain both distance and orientations information, instead of evaluating them separately, we directly use the TM-score to measure the accuracy of the predicted 3D structures obtained using these outputs information as the constraints in trRosetta folding protocol. In Table 4, we list the performance of OPUS-Contact and trRosetta ^5^ on CAMEO-Hard61 (60), CAMEO (78), CASP13 (26) and CASP14 (15). Both OPUS-Contact and trRosetta use the same multiple sequence alignment results from DeepMSA ^37^.

**Table 4.**
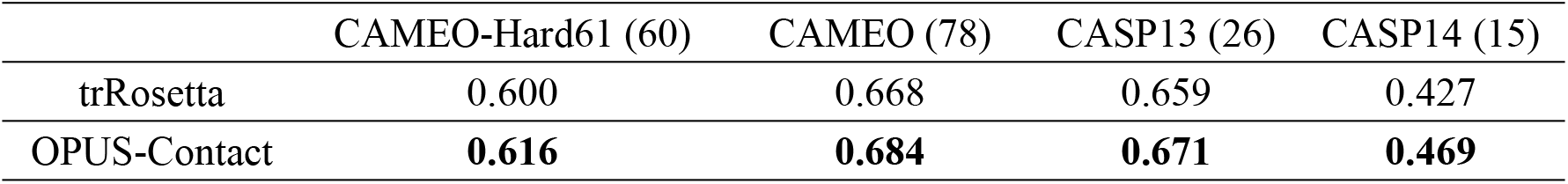
TM-score of OPUS-Contact and trRosetta on different datasets.

### Performance of OPUS-Fold2

We compare the folding performance of OPUS-Fold2 and the Rosetta ^22, 23^ folding protocol in trRosetta ^5^ on CAMEO-Hard61 (60). As shown in Figure 5, when using the distance constraints exclusively, OPUS-Fold2 outperforms trRosetta by a large margin. OPUS-Fold2 also slightly outperforms trRosetta when using both distance and orientations constraints. However, the complete folding protocol of trRosetta includes some other terms that haven’t been included into OPUS-Fold2 yet. The final result of the complete version of trRosetta is slightly better than that of OPUS-Fold2.

**Figure 5.**
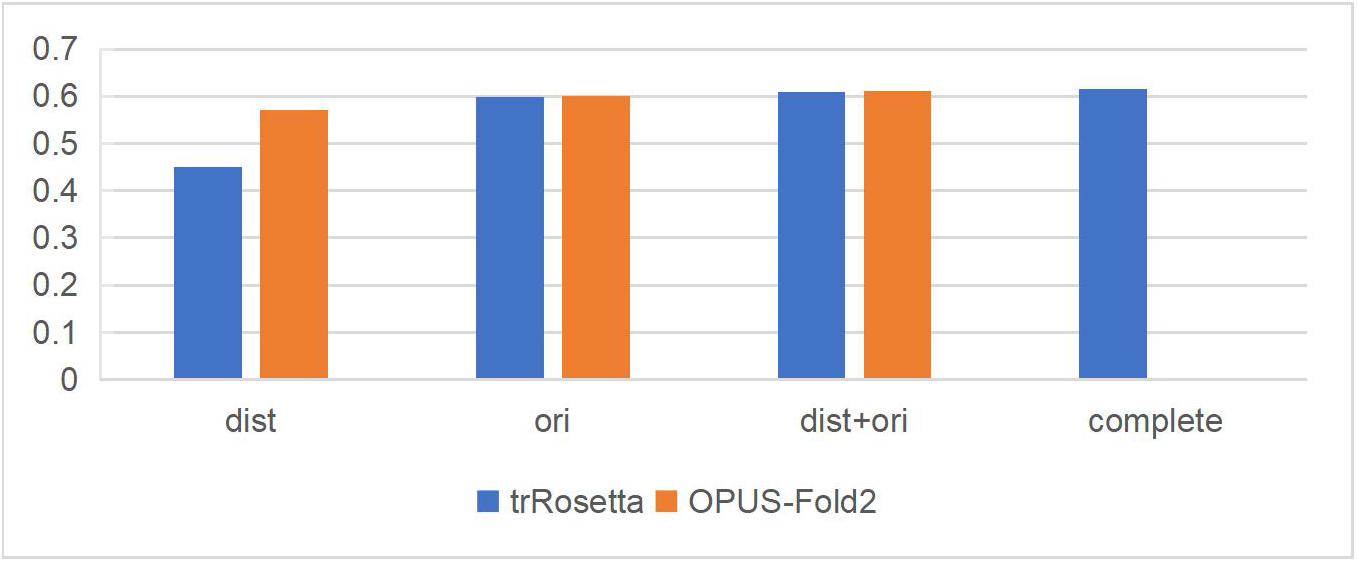
Performance of OPUS-Fold2 and the Rosetta folding protocol in trRosetta based on the outputs from OPUS-Contact on CAMEO-Hard61 (60). *dist* denotes the prediction obtained by distance-guided folding exclusively, *ori* denotes the prediction obtained by orientations-guided (ω, θ and φ) folding exclusively, *dist+ori* denotes the prediction obtained using both of them, and *complete* denotes the prediction obtained using trRosetta’s original complete energy terms (including the ramachandran, the omega, the van der Waals (vdw), and the centroid backbone hydrogen bonding (cen_hb) terms). The y-axis represents the TM-score.

We list the results of OPUS-Fold2 and the results of the complete version of trRosetta in Figure 6. OPUS-Fold2 exhibits a consistent well performance on all four datasets and achieves comparable performance to trRosetta when using identical inputs from OPUS-Contact.

**Figure 6.**
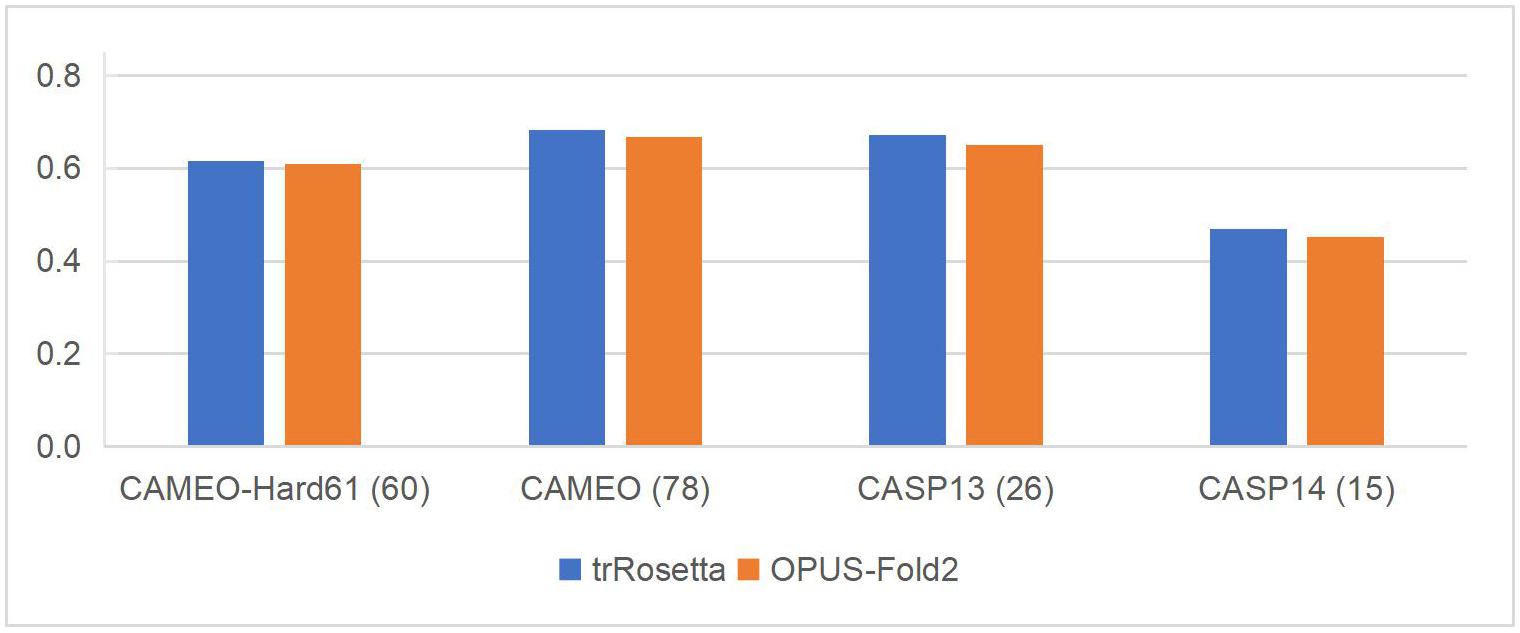
Performance of OPUS-Fold2 and the complete version of trRosetta on CAMEO-Hard61 (60), CAMEO (78), CASP13 (26) and CASP14 (15).

We show the optimization process of OPUS-Fold2 in Figure 7. The total loss become lower and the TM-score become higher along with the optimization. We also show some intermediate structures during the optimization process of OPUS-Fold2 in Figure 8, Figure 9 and Figure 10. For example, Figure 8 shows the optimization for target *2020-01-18_00000081_1.pdb* (with 444 residues in length) in CAMEO-Hard61 (60). In the first 100 epochs, the loss rapidly decreases from −67 to −130, and the TM-score rapidly increases from 0.247 to 0.792. The loss continually decreases in the following epochs and stabilizes around −140, and the TM-score stabilizes around 0.87.

**Figure 7.**
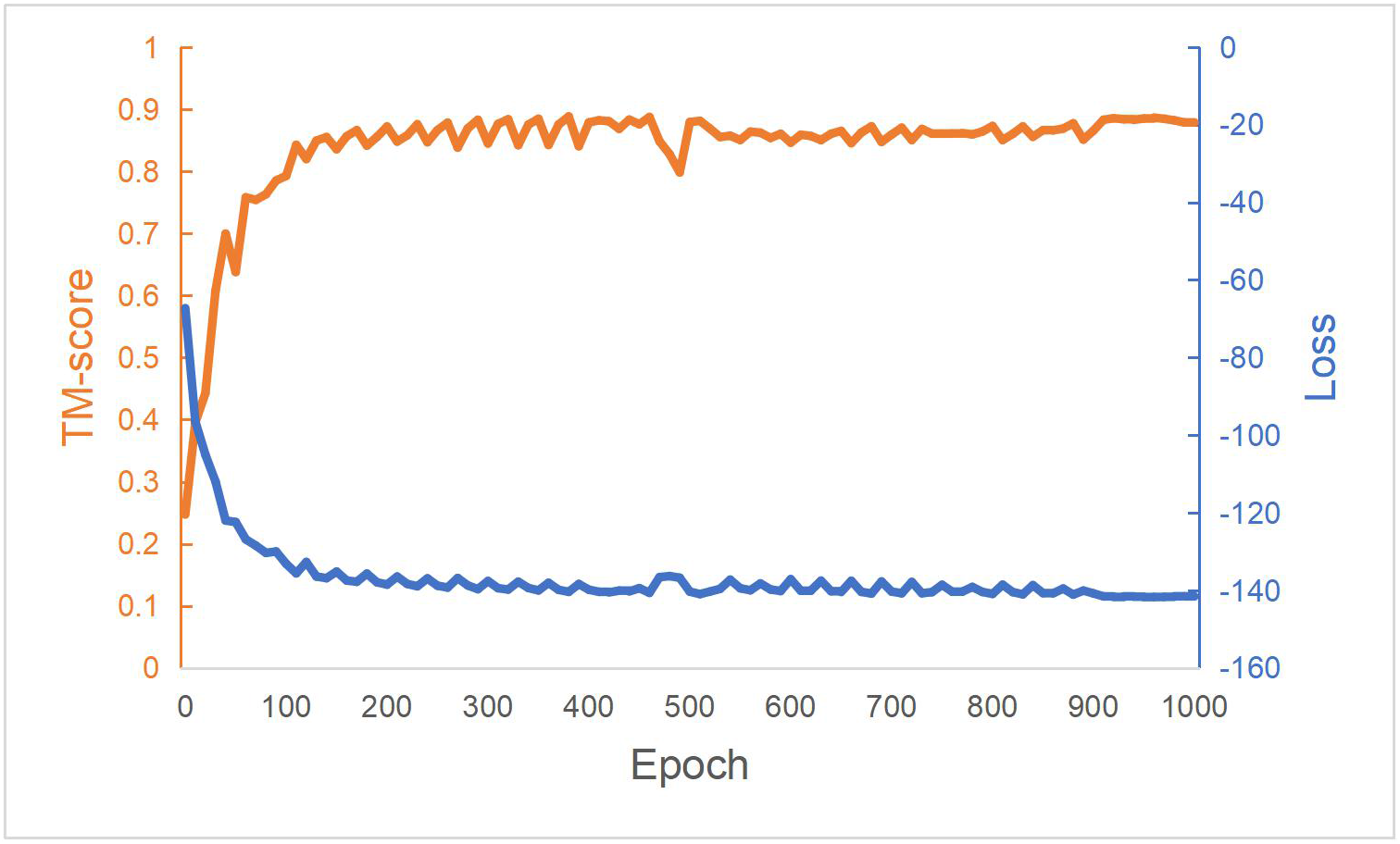
OPUS-Fold2 optimization process of target *2020-01-18_00000081_1.pdb* (with 444 residues in length) in CAMEO-Hard61 (60). The blue line is the total loss and the red line is the TM-score.

**Figure 8.**
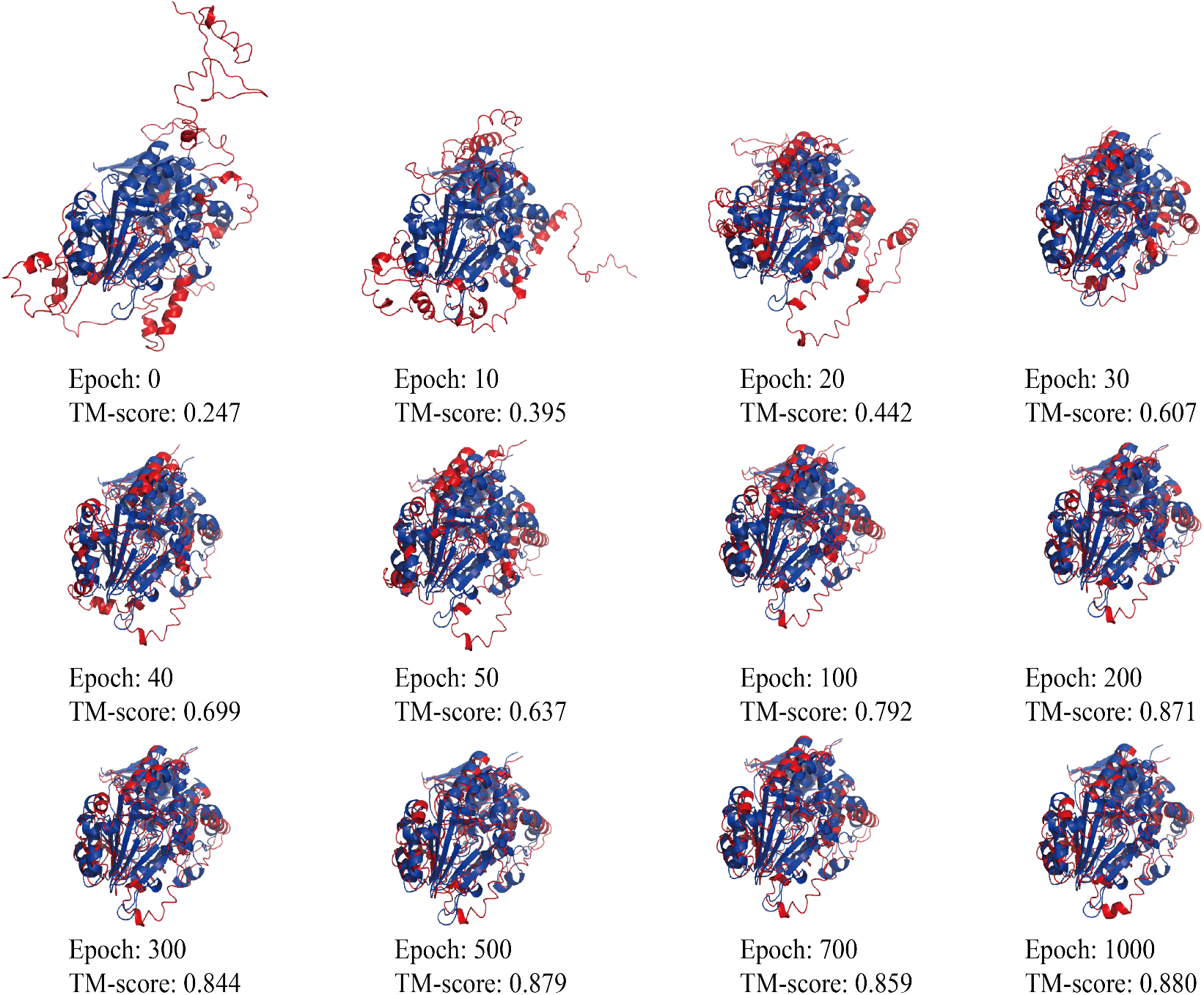
Some intermediate structures of target *2020-01-18_00000081_1.pdb* (with 444 residues in length) during the optimization process of OPUS-Fold2.

**Figure 9.**
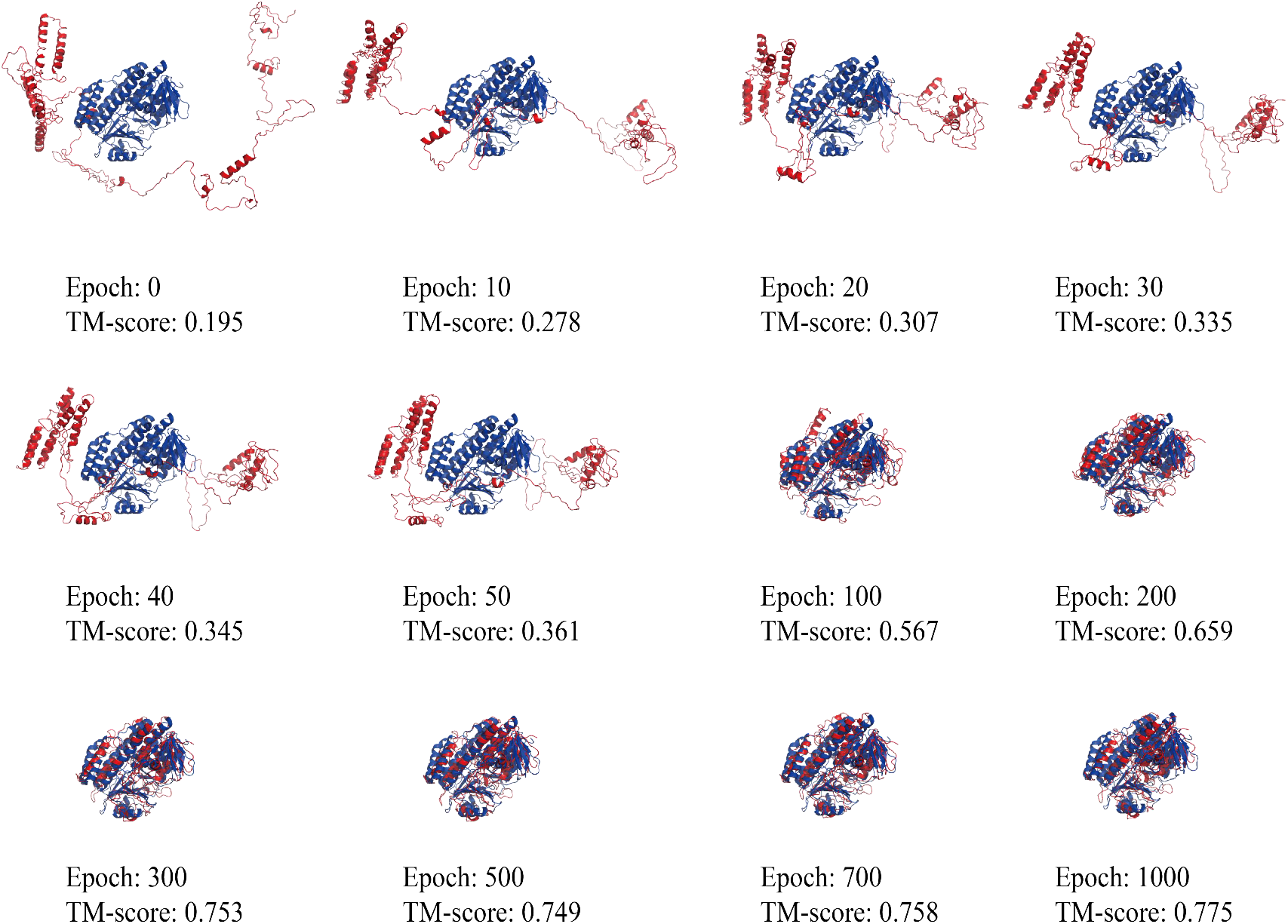
Some intermediate structures of target *6BZT_D_21_522.pdb* (with 501 residues in length) during the optimization process of OPUS-Fold2.

**Figure 10.**
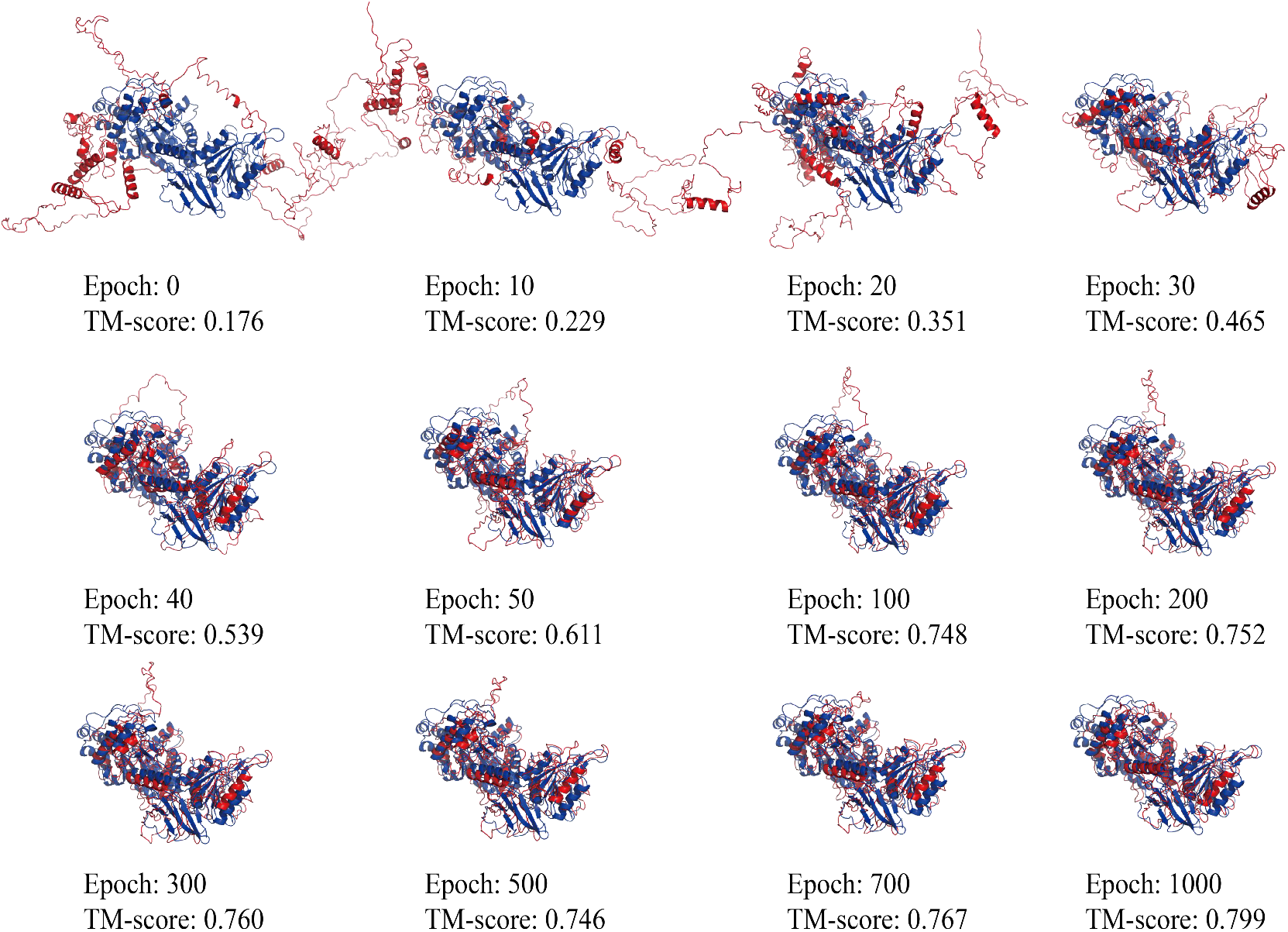
Some intermediate structures of target *2020-02-08_00000106_1.pdb* (with 665 residues in length) during the optimization process of OPUS-Fold2.

## Concluding Discussion

Protein 3D structure prediction is an important and challenging task. The feasibility of it has been demonstrated by the AlphaFold2 in CASP14 ^7^. In this paper, we develop an open-source toolkit for protein 3D structure modeling, named OPUS-X. It includes a state-of-the-art protein torsion angles, secondary structure and solvent accessibility predictor, namely OPUS-TASS2; a better global distance and orientations constraints predictor compared with the open-source state-of-the-art method trRosetta ^5^, namely OPUS-Contact; and a gradient-based protein folding framework that is comparable to the Rosetta ^22, 23^ folding protocol in trRosetta ^5^, namely OPUS-Fold2.

As shown in Table 1, OPUS-TASS2 outperforms NetSurfP-2.0 ^12^, SPOT-1D ^9^ and OPUS-TASS ^10^ by a large margin, especially on the most difficult dataset CASP14 (15). We believe the accurate and detailed distance and orientations global information plays a dominant role. Table 2 also indicates the importance of global information. To further demonstrate the importance of global information and the potentiality of OPUS-TASS2, we feed the real values of global information from the native structures into OPUS-TASS2 to predict their 1D features. The results (Table 3) show that, using the real orientation information (ω, θ and φ), real distance information, and both of them will significantly improve protein 1D features prediction accuracy. Note that, the improvement of introducing real orientations information is significantly larger than that of introducing real distance information, indicating the dominant influence of global orientations information. Combining them will further boost the final accuracy.

Since the trRosetta-style’s ^5^ outputs contain both distance and orientations information, and they may need to achieve a trade-off to deliver better 3D structure prediction, evaluating them separately may not be a good idea. For example, as shown in Figure 4, traditional distance-based metrics are not significant correlated with the final 3D prediction accuracy. Therefore, we directly use the final 3D prediction results to evaluate the trRosetta-style’s outputs. Comparing with the open-source state-of-the-art method trRosetta, OPUS-Contact achieves better 3D structure prediction accuracy on CAMEO-Hard61 (60), CAMEO (78), CASP13 (26) and CASP14 (15) (Table 4).

We also compare the performance of OPUS-Contact with that of some other methods on CASP13 (26) and CASP14 (15). The results of their methods are downloaded from the CASP website. On CASP13 (26), both OPUS-Contact (TM-score=0.671) and trRosetta (TM-score=0.659) are better than the best method at the time A7D (TM-score=0.644) ^1^. On CASP14 (15), both OPUS-Contact (TM-score=0.469) and trRosetta (TM-score=0.427) are lower than the best human group method AlphaFold2 (TM-score=0.850) and the best server group method Zhang-Server (TM-score=0.540) ^6^. We believe the reason may lies in the insufficient multiple sequence alignment searching step since the alignment results of 5 out of 15 targets have less than 5 sequences. Nevertheless, comparing with the other methods, OPUS-Contact provides a better open-source protein structure prediction tool that can be run on the user’s own server for the community.

OPUS-Fold2 is a gradient-based protein folding method. It is written in Python and TensorFlow2.4, easily to be modified at source-code level, which is especially useful for the folding energy term developers. Figure 5 shows the contributions of distance and orientations constraints. Same as the Rosetta ^22, 23^ folding protocol in trRosetta ^5^, the folding results guided by orientations constraints are significantly better than that guided by distance constraints. After combining them together, the accuracy is further improved. On CAMEO-Hard61 (60), OPUS-Fold2 outperforms trRosetta when using distance constraints exclusively, orientations constraints exclusively and both of them jointly as the energy function. However, after introducing some other terms such as the ramachandran, the omega, the van der Waals, and the centroid backbone hydrogen bonding into the trRosetta’s energy function, the folding performance of trRosetta is slightly better than that of OPUS-Fold2 on CAMEO-Hard61 (60), CAMEO (78), CASP13 (26) and CASP14 (15) (Figure 6). One of our future goals is to add these terms into OPUS-Fold2. Figure 7 and Figure 8 show some insights of the OPUS-Fold2 optimization step. Along with the optimization, the total loss descends logically, indicating the effectiveness of OPUS-Fold2.

## Acknowledgements

The work was partially supported by Shanghai Municipal Science and Technology Major Project (No.2018SHZDZX01), and ZJLab. QW thanks the Welch Foundation (Q-1826) for support. JM thanks the support from the Welch Foundation (Q-1512).

## Author contributions

Gang Xu and Jinapeng Ma designed the project. Gang Xu conducted the experiments. All authors contributed to the manuscript composing.

## TOC graphic

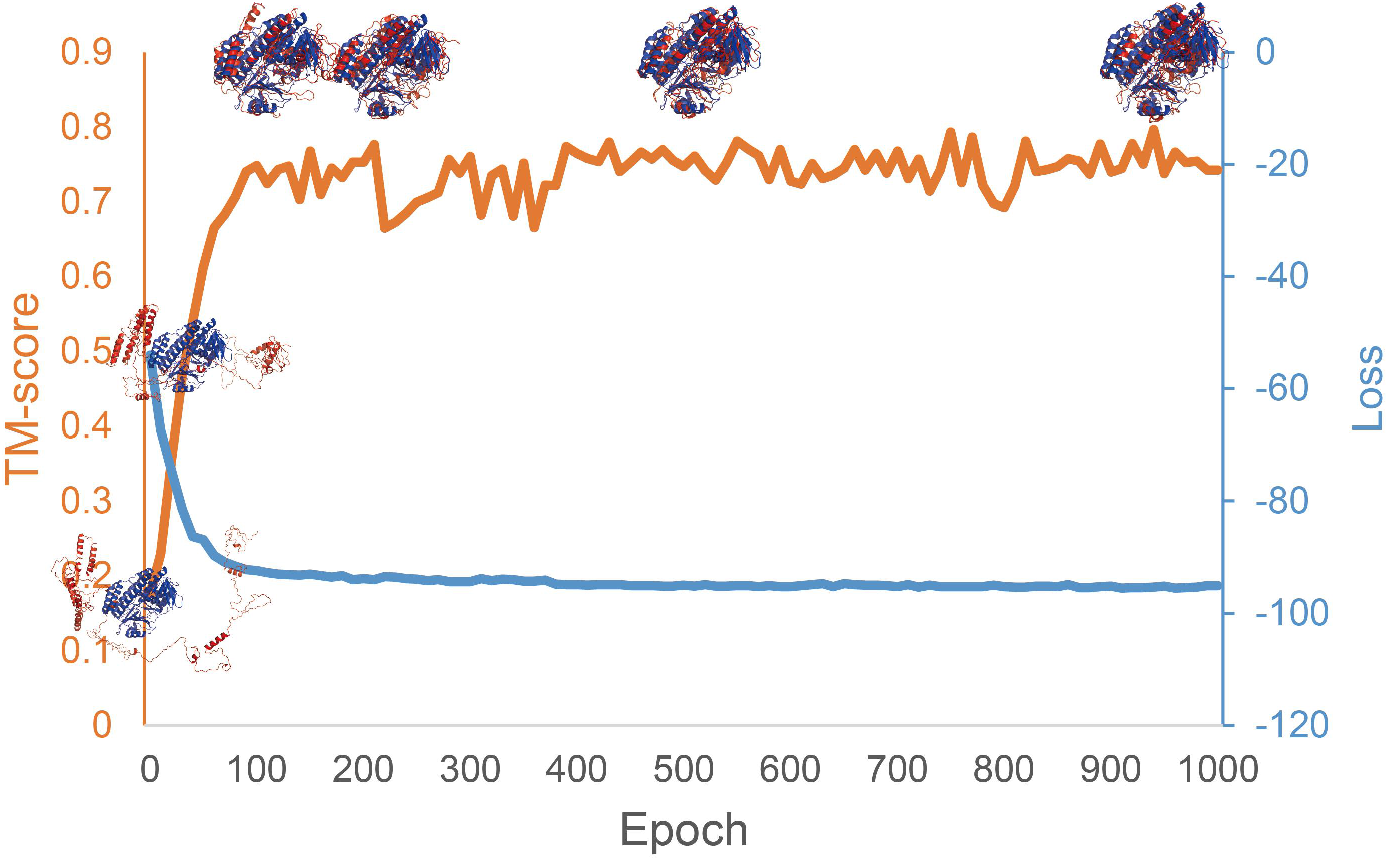

